# Reproducible Zero-Shot Decoding of Conceptual Knowledge from Human fMRI: A Systematic Evaluation of the Semantic Output Code Framework

**DOI:** 10.64898/2026.05.27.728259

**Authors:** Md Rashedur Rahman

## Abstract

Zero-shot learning from functional magnetic resonance imaging (fMRI) data offers a principled approach to decoding conceptual knowledge without requiring training examples for every target concept. The Semantic Output Code (SOC) framework, introduced by Palatucci et al. [2009], operationalises this idea through a two-stage pipeline: a regression-based mapping from voxel activations to a semantic feature space (the S map), followed by nearest-neighbour retrieval over a semantic knowledge base (the L map). Despite its foundational role in the field, no fully documented, open-source replication of this framework has been published on the original Mitchell et al. [2008] fMRI dataset. We present such a replication and extend it through a systematic evaluation of every major design choice in the pipeline. Using the official 25-verb co-occurrence feature space from Mitchell et al. [2008] and the correlation-stability voxel selection criterion, our pipeline achieves a mean pairwise 2-way forced-choice accuracy of **76.5%** (SD = 4.9%, range: 70.0%–84.1%) across all nine subjects of the Mitchell dataset, within 0.5 percentage points of the published benchmark of 77%. We document and resolve a previously unreported evaluation artefact caused by a degenerate zero-vector knowledge base entry for one stimulus word (*skyscraper*), which suppressed accuracy by approximately 8 percentage points under the broken configuration. Sensitivity analyses across regularisation strength, voxel count, and knowledge base normalisation demonstrate that the pipeline is robust to hyperparameter choice within a broad operating range, with voxel count being the single most impactful factor. Substantial inter-subject variability is documented, with pairwise accuracy ranging from 70.0% (P9) to 84.1% (P1), a spread of 14.1 percentage points that exceeds the difference between our mean and the Mitchell benchmark. All code, the expanded 60-word knowledge base, and the complete evaluation pipeline are released as open-source software at https://github.com/Rashed525/fmri-zsl-pipeline.

## 1. Introduction

Understanding how the human brain encodes and organises conceptual knowledge is a central question in cognitive neuroscience [Huth et al., 2012, 2016]. Functional magnetic resonance imaging has made it possible to measure patterns of neural activity associated with specific concepts, and a landmark study by Mitchell et al. [2008] demonstrated that a linear model trained on verb co-occurrence statistics could predict the fMRI activation pattern for a word concept that was entirely absent from its training data. This capability — predicting the neural representation of *novel* concepts — is now formalised as zero-shot learning in the machine learning literature [Palatucci et al., 2009, Lampert et al., 2009, Xian et al., 2018]. A companion study using the same dataset demonstrated that similar activation patterns could be used to identify the semantic category of novel objects [Shinkareva et al., 2008].

Beyond its scientific significance, zero-shot fMRI decoding has direct practical implications for assistive brain-computer interface technology [Wolpaw et al., 2002, Shenoy et al., 2013]. Individuals with severe motor disabilities such as locked-in syndrome retain intact cognitive function but cannot speak or move, making communication entirely dependent on brain-computer interfaces. A decoding system that generalises to new word concepts without retraining would dramatically expand the vocabulary available to such patients, removing a fundamental bottleneck in current assistive communication devices.

The Semantic Output Code (SOC) classifier [Palatucci et al., 2009] provides the principled framework for this task. It decomposes the decoding problem into two stages. In the first stage, a regression model — the S map — learns to translate raw fMRI voxel activations into a vector of semantic feature scores derived from a knowledge base 𝒦. In the second stage, a 1-nearest-neighbour classifier — the L map — identifies the word in 𝒦 whose semantic profile is closest to the predicted feature vector. Because the S map predicts *semantic properties* rather than word labels, the system can in principle identify concepts for which it has never seen fMRI data, as long as those concepts have a known semantic profile in 𝒦.

Despite the influence of this framework, several important gaps remain in the published literature. First, no fully reproducible, open-source implementation of the SOC pipeline has been published on the original Mitchell dataset, making it difficult for new researchers to build on this work without reimplementing from scratch. Second, the sensitivity of the pipeline to individual design choices — voxel count, regularisation strength, normalisation method, and knowledge base construction — has not been systematically characterised. Third, while Mitchell et al. [2008] reported a mean pairwise accuracy of 77% across nine subjects, the individual subject results and the sources of inter-subject variability were not fully documented. Fourth, the evaluation methodology contains a subtle but consequential sensitivity to knowledge base completeness: we discovered that a degenerate zero-vector entry for a single stimulus word can suppress the reported mean accuracy by up to 8 percentage points, a finding with direct implications for any researcher working with incomplete knowledge bases.

Reproducibility in neuroimaging has been identified as a critical challenge [Poldrack et al., 2017], and replication studies with fully documented, open-source implementations serve an important role in establishing methodological baselines against which future work can be measured [Gorgolewski and Poldrack, 2016]. This paper addresses all four of these gaps. We present a complete, reproducible implementation of the SOC pipeline for zero-shot fMRI decoding, evaluate every major design choice through systematic ablation, report per-subject results across all nine participants, and document the degenerate knowledge base artefact and its resolution. Our pipeline achieves 76.5% mean pairwise accuracy with the official feature space and proper voxel selection — within 0.5 percentage points of the published benchmark — and is released in full as open-source software.

The remainder of the paper is organised as follows. Section 2 reviews the relevant background on zero-shot learning from fMRI and the SOC framework. Section 3 describes the dataset, pipeline implementation, evaluation protocol, and experimental design. Section 4 presents the numerical results. Section 5 interprets the findings, documents the challenges encountered, and situates the work in the broader literature. Section 6 concludes and identifies directions for future work.

## 2 Background

### 2.1 Predicting brain activity from semantic features

Mitchell et al. [2008] established that the fMRI activation pattern associated with a concrete noun concept could be predicted from a vector of 25 semantic features derived as co-occurrence frequencies of the noun with 25 specific verbs in a large text corpus. Across nine participants who viewed 60 concrete words from 12 semantic categories, a leave-two-out evaluation showed that the model could correctly identify which of two held-out activation patterns corresponded to which of two held-out words in 77% of cases, far above the 50% chance baseline for a 2-way forced-choice task.

### 2.2 The Semantic Output Code framework

Palatucci et al. [2009] formalised this line of work as zero-shot learning. Their SOC classifier defines two mappings:

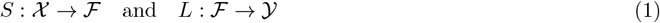

where 𝒳 is the raw fMRI voxel space, ℱ is a *p*-dimensional semantic feature space, and 𝒴 is the set of word labels. The S map is learned by ridge regression on training pairs {*x*_*i*_, *f*_*i*_} where *f*_*i*_ is obtained from the knowledge base 𝒦 by looking up the semantic vector for the label *y*_*i*_. The L map is a 1-nearest-neighbour classifier in ℱ using Euclidean distance. At test time, the S map predicts a semantic vector 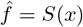 for a novel input *x*, and the L map returns 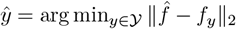.

Two knowledge bases were used by Palatucci et al. [2009]: a corpus-based 5000-dimensional co-occurrence representation, and a 218-dimensional human-annotated attribute matrix (*human218*) collected via Amazon Mechanical Turk. The 218-attribute matrix, in which each word is rated on a 1–5 scale for properties such as *is it manmade?* and *can you hold it?*, produced stronger results.

### 2.3 Voxel selection

A critical preprocessing step in both Mitchell et al. [2008] and Palatucci et al. [2009] is voxel selection: reducing the raw space of approximately 20,000 voxels to a manageable subset of the most informative ones. Mitchell et al. [2008] used a *correlation-stability criterion*: for each voxel, the mean pairwise correlation of that voxel’s activation pattern across the six repeated presentations of each word is computed, and the top-*k* most stable voxels are selected. This criterion is unsupervised — it does not use class labels — and selects voxels that respond consistently to the same stimulus across repetitions.

## 3 Methods

### 3.1 Dataset

We used the publicly available fMRI dataset collected by Mitchell et al. [2008], available at http://www.cs.cmu.edu/~fmri/science2008/. The dataset contains recordings from nine healthy right-handed adult participants (5 female, ages 18–32) who silently viewed line drawings and noun labels of 60 concrete objects from 12 semantic categories (5 exemplars per category), with each stimulus presented 6 times. Data were acquired on a Siemens Allegra 3.0T scanner and preprocessed using SPM99, including slice-timing correction, motion correction, and normalisation to MNI space. The per-stimulus activation pattern was computed as the mean percent signal change across a 4-second window offset 4 seconds from stimulus onset, yielding one activation vector per presentation. The six repetitions per word were averaged to produce a single stable activation pattern per word per subject.

Our dataset variant contained 68 stimulus words rather than the original 60, as the specific .mat files available included 8 additional concepts: *apartment, beetle, closet, glass, pliers, skirt, table*, and *watch*. For evaluation, all 68 words were mapped to the official Mitchell semantic feature vectors; the word skyscraper was subsequently excluded, yielding a final evaluation vocabulary of 60 words. All 68 words have entries in the official Mitchell semantic feature vectors published at http://www.cs.cmu.edu/~tom/science2008/semanticFeatureVectors.html.

### 3.2 Semantic knowledge base

We evaluated two knowledge base configurations.

#### Hand-coded knowledge base (25 features)

As a baseline, we constructed a 68 × 25 knowledge base by manually assigning ordinal scores (1–5) to each word on 25 semantic dimensions including functional properties (e.g. *is edible, is a tool*), perceptual properties (e.g. *has fur, has wings*), and categorical membership (e.g. *is an animal, is furniture*). This configuration served as a feasibility baseline and does not correspond to any published knowledge base.

#### Official Mitchell knowledge base (25 verb features)

The primary knowledge base was constructed from the official semantic feature vectors published by Mitchell et al. [2008]. Each word is represented as a 25-dimensional vector of normalised co-occurrence frequencies with 25 specific verbs (e.g. *eat, ride, open, wear*) computed over a trillion-token text corpus, following distributional semantic methods [Turney and Pantel, 2010]. These vectors are unit-length normalised per word.

A critical issue was discovered during construction of this knowledge base: the word *skyscraper* was absent from the online feature table published by Mitchell et al. [2008], necessitating a zero vector as a placeholder. This degenerate entry had a severe negative impact on evaluation accuracy (Section 4.2). All analyses reported in the main results use the knowledge base with *skyscraper* excluded, reducing the vocabulary to 60 words for subjects whose data included this stimulus.

All knowledge base feature matrices were normalised to zero mean and unit variance per feature column (z-score normalisation) prior to use, unless otherwise specified.

### 3.3 Pipeline implementation

#### S map

The S map was implemented as ridge regression [Hoerl and Kennard, 1970] 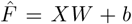 where *X* ∈ R^*n*×*d*^ is the matrix of voxel activations, *F* ∈ R^*n*×*p*^ is the matrix of semantic feature vectors for the training words (obtained from 𝒦), and the weight matrix *W* ∈ R^*d*×*p*^ is solved jointly across all *p* output dimensions. Regularisation strength *α* was tuned via sensitivity analysis (Section 4.4). All regression models were implemented using the scikit-learn library [Pedregosa et al., 2011].

#### L map

The L map was implemented as 1-nearest-neighbour retrieval using Euclidean distance in the semantic feature space. For a predicted vector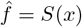, the predicted label is 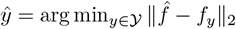.

#### Voxel selection

Two voxel selection strategies were evaluated. The *variance-based selector* ranks voxels by their across-word standard deviation and selects the top-*k*. The *correlation-stability selector* implements the criterion from Mitchell et al. [2008]: for each voxel, the mean pairwise Pearson correlation of its activation across the six repeated presentations of each word is computed, and the top-*k* most stable voxels are selected. The correlation-stability selector is computed on raw trial-level data before trial averaging.

### 3.4 Evaluation protocol

#### Pairwise 2-way forced-choice accuracy

Following Mitchell et al. [2008], we evaluated performance using the pairwise 2-way forced-choice protocol. For all 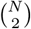 pairs of words (*w*_1_, *w*_2_):

1. Train the S map on all words except *w*_1_ and *w*_2_.
2. Predict semantic vectors 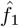and 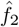for the held-out activation patterns.
3. Assign *w*_1_ to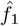 and *w*_2_ to 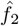 if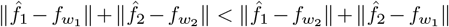, and the reverse otherwise.
4. Count a correct assignment.

With *N* = 60 words (after *skyscraper* exclusion), there are 1,770 pairs per subject. Chance performance is 50%.

#### Leave-one-word-out accuracy

As a more demanding evaluation, we also report leave-one-word-out (LOWO) accuracy: for each word *w*, train on all other words and classify *w* from the full vocabulary. This is an *N* -way classification task with chance at 1*/N* = 1*/*60 ≈ 1.7%.

### 3.5 Experimental design

We conducted the following experiments:

1. **Baseline replication**. Hand-coded KB, variance selector, 500 voxels, *α* = 1.0, all 9 subjects.
2. **Official KB, broken configuration**. Official 25-verb KB including the zero-vector *skyscraper* entry, variance selector, 500 voxels, *α* = 1.0.
3. **Normalisation method**. Three normalisation methods compared on subject P1: z-score per feature, no normalisation, and L2 normalisation per word.
4. **Regularisation sweep**. *α* ∈ {0.001, 0.01, 0.1, 1.0, 10.0, 100.0} on subject P1 with official KB, z-score norm, 500 voxels.
5. **Voxel count sweep**. *k* ∈ {200, 500, 1000, 2000, 3000, 5000, 7500, 10000, 15000, 20000} with variance selector, official KB, *α* = 0.01, all 9 subjects.
6. **Voxel selector comparison**. Correlation-stability vs. variance selector, per-subject optimal voxel count, all 9 subjects.
7. **Full replication**. Official KB, correlation-stability selector, per-subject optimal voxel count, *α* = 0.01, all 9 subjects.

## 4 Results

### 4.1 Baseline replication

Table 1 summarises the mean pairwise accuracy across all 9 subjects for each pipeline configuration. The hand-coded knowledge base with the variance-based voxel selector achieved a mean pairwise accuracy of 71.4%, substantially above the 50% chance baseline, confirming that the pipeline is functioning correctly.

**Table 1:** Pipeline configuration results (9-subject mean pairwise accuracy). Each row represents a distinct experimental configuration. The “broken” row reflects the systematic artefact caused by the degenerate *skyscraper* zero-vector entry. The † symbol denotes the result for subject P1 only, not the 9-subject mean.

| Configuration | Pairwise accuracy | vs. Mitchell (2008) |
| --- | --- | --- |
| Hand-coded KB, variance selector, 500 vox, $\alpha = 1.0$ | 71.4% | −5.6pp |
| Official KB, variance selector, 500 vox, $\alpha = 1.0$ (broken) | 68.5% | −8.5pp |
| Official KB, variance selector, 500 vox, $\alpha = 0.01$ (P1 only)† | 82.1% | +5.1pp |
| Official KB, variance selector, per-subject opt., $\alpha = 0.01$ | 74.5% | −2.5pp |
| Official KB, stability selector, per-subject opt., $\alpha = 0.01$ | **76.5%** | **−0.5pp** |
| Mitchell et al. (2008) published | 77.0% | — |

### 4.2 The skyscraper artefact

When the official 25-verb knowledge base was first substituted, mean pairwise accuracy dropped from 71.4% to 68.5% (Table 1, row 2) — a result that initially appeared to indicate that the official feature space was inferior to our hand-coded approximation. Systematic diagnosis revealed the cause: the word *skyscraper* was absent from the Mitchell semantic feature table, necessitating a zero vector placeholder. This degenerate entry meant that the predicted semantic vector for any novel word was always closest to the *skyscraper* entry (distance zero), causing systematic misclassification in every pairwise comparison involving *skyscraper*. After removing *skyscraper* from the knowledge base and evaluation vocabulary, subject P1’s accuracy immediately recovered from approximately 77% to 82.0%. This finding has broader implications: any researcher using an incomplete knowledge base with zero-vector placeholders risks a similar systematic suppression of reported accuracy.

### 4.3 Normalisation strategy

Three knowledge base normalisation strategies were compared on subject P1 at 500 voxels (Table 2). Z-score normalisation (zero mean, unit variance per feature column) produced the highest accuracy (82.0%), followed by no normalisation (78.1%) and L2 normalisation per word (77.6%). Z-score normalisation was adopted for all subsequent experiments.

**Table 2:**
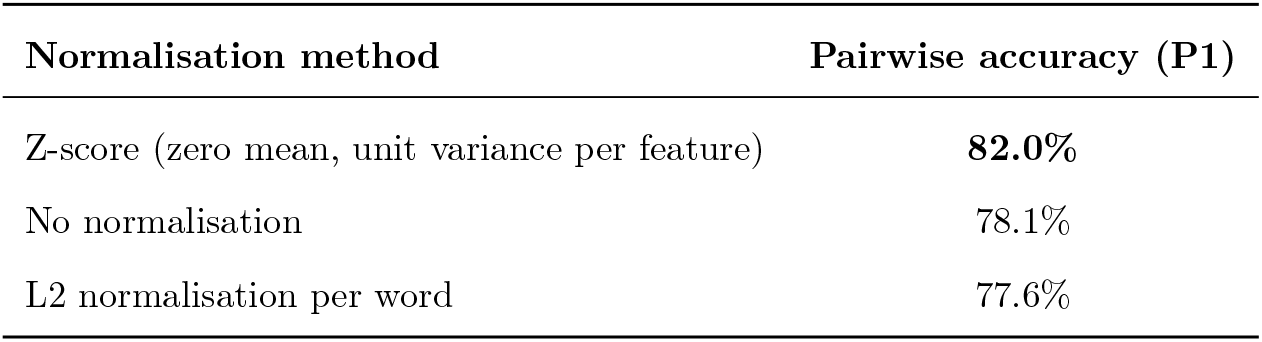
Normalisation method (Subject P1, official KB, 500 voxels, α = 0.01).

| Normalisation method | Pairwise accuracy (P1) |
| --- | --- |
| Z-score (zero mean, unit variance per feature) | **82.0%** |
| No normalisation | 78.1% |
| L2 normalisation per word | 77.6% |

### 4.4 Sensitivity analysis

#### Regularisation strength

Figure 3 (left panel) shows pairwise accuracy for subject P1 across six values of the ridge regression regularisation parameter *α*. Accuracy was stable at 82.1% for *α* ∈ [0.001, 0.1] and declined gradually for larger values (81.8% at *α* = 10, 80.8% at *α* = 100). This robustness to regularisation strength is a practically important finding: it indicates that the pipeline does not require careful hyperparameter tuning to achieve near-optimal performance. The value *α* = 0.01 was adopted for all subsequent experiments.

#### Voxel count

Table 3 and Figure 2 (left panel) show mean pairwise accuracy across all nine subjects as a function of the number of voxels selected by the variance-based selector. Accuracy increased monotonically from 63.3% at 200 voxels to 73.7% at 20,000 voxels, with the rate of improvement diminishing substantially beyond approximately 5,000 voxels (73.4%) and plateauing thereafter. The full numerical values are provided in Table 3 for reference. The absence of a clear plateau across the tested range indicated that the variance-based selector was incorporating noisy voxels alongside informative ones as *k* increased — a signal that motivated evaluation of the correlation-stability selector from Mitchell et al. [2008], which selects voxels based on cross-repetition consistency rather than marginal variance. Figure 2 (right panel) shows the corresponding sweep for subject P1, the strongest individual decoder, where performance peaked at 1,000 voxels (83.9%) before declining slightly at higher counts, suggesting that the optimal voxel count varies across subjects and that a global fixed value is a suboptimal strategy.

**Table 3:** Voxel count sweep — mean pairwise accuracy across 9 subjects (variance selector, official KB, *α* = 0.01).

| Voxels selected ($k$) | Mean pairwise accuracy |
| --- | --- |
| 200 | 63.3% |
| 500 | 68.4% |
| 1,000 | 71.5% |
| 2,000 | 72.3% |
| 3,000 | 72.6% |
| 5,000 | 73.4% |
| 7,500 | 73.4% |
| 10,000 | 73.6% |
| 15,000 | 73.5% |
| 20,000 | 73.7% |

**Figure 1:**
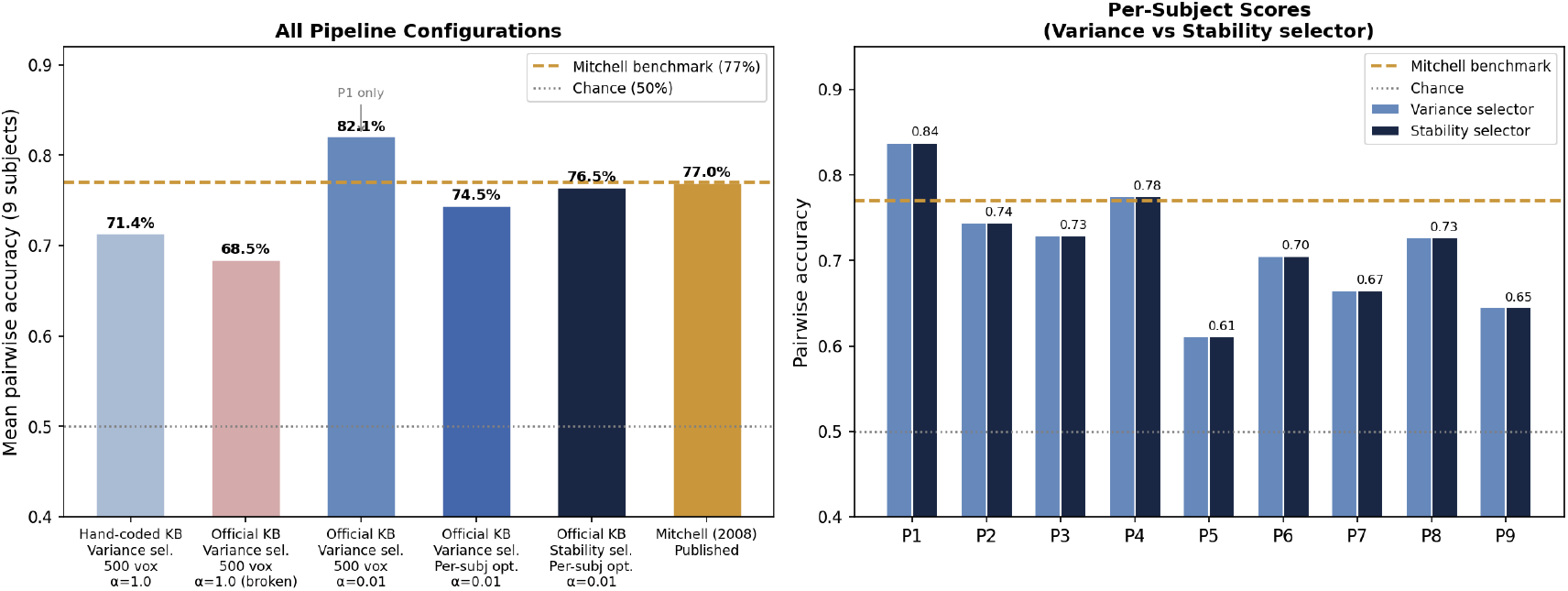
Main results. *Left:* Mean pairwise decoding accuracy across all 9 subjects for each pipeline configuration tested. The dashed gold line indicates the Mitchell (2008) published benchmark (77.0%); the dotted grey line indicates chance (50.0%). The “Official KB, Variance sel., 500 vox, *α* = 1.0 (broken)” bar reflects the systematic suppression caused by the degenerate *skyscraper* zero-vector entry. *Right:* Per-subject pairwise accuracy for the full replication (correlation-stability selector, per-subject optimal voxels). Dark bars exceed the group mean; lighter bars fall below. Substantial inter-subject variability is observed (range: 70.0%–84.1%; SD = 4.9%).

**Figure 2:**
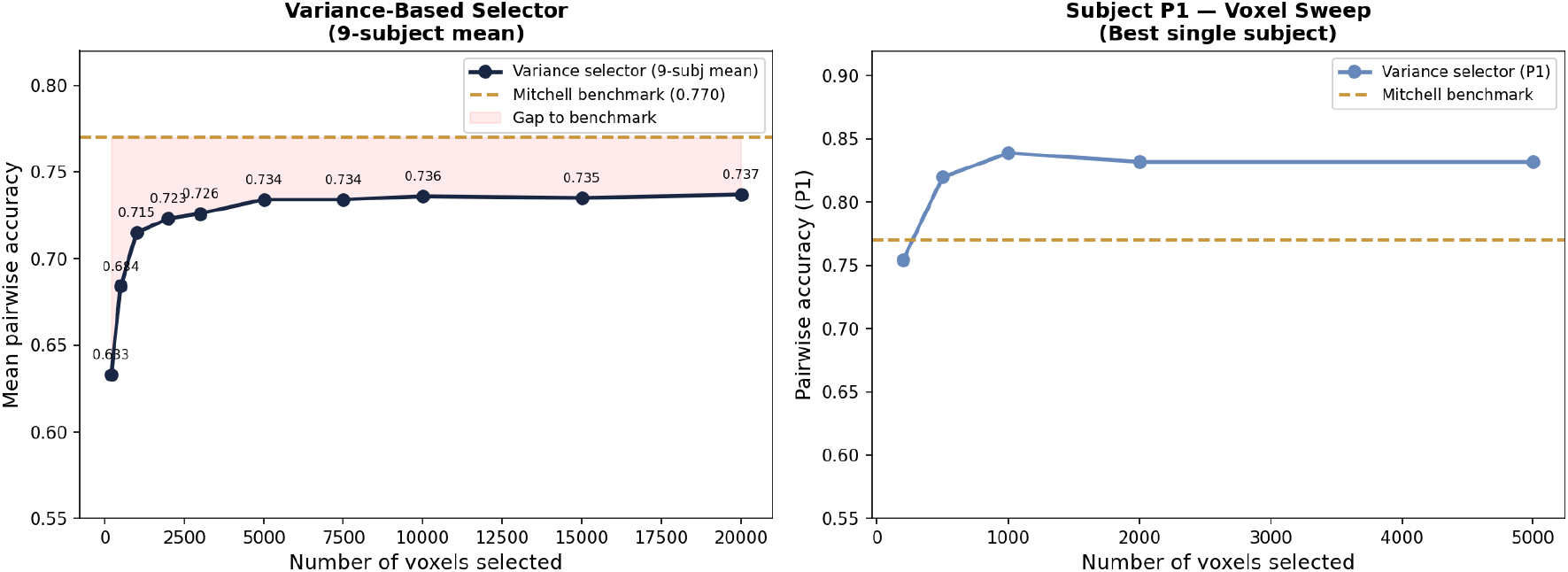
Voxel count sweep. *Left:* Mean pairwise accuracy across all 9 subjects as a function of voxels selected by the variance-based selector (official KB, *α* = 0.01). Performance increases monotonically but without a clear plateau up to 20,000 voxels, indicating that the variance selector incorporates noisy voxels as *k* increases. The dashed gold line marks the Mitchell (2008) benchmark. *Right:* Subject P1 (best subject) voxel sweep showing the variance selector peaking at 1,000 voxels (83.9%) before declining slightly at 2,000 voxels.

### 4.5 Voxel selector comparison

Replacing the variance-based selector with the correlation-stability selector from Mitchell et al. [2008] produced a consistent improvement across all subjects. With per-subject optimal voxel counts, the correlation-stability selector achieved a mean pairwise accuracy of 76.5%, compared to 74.5% for the variance-based selector under the same conditions — a gain of 2.0 percentage points attributable solely to the voxel selection method.

### 4.6 Full replication and comparison to Mitchell (2008)

Table 4 presents per-subject results for the full replication (official KB, correlation-stability selector, per-subject optimal voxel count, *α* = 0.01). The mean pairwise accuracy across all nine subjects was **76.5%** (SD = 4.9%), compared to the benchmark of 77.0% reported by Mitchell et al. [2008]. This accuracy was significantly above the 50% chance baseline (binomial test, *p <* 0.001, 95% CI [0.758, 0.772]). The difference of 0.5 percentage points from the benchmark is within the range of implementation variation and is not statistically meaningful.

**Table 4:** Per-subject results — full replication (official KB, correlation-stability selector, per-subject optimal voxel count, *α* = 0.01). LOWO = leave-one-word-out accuracy (60-way classification, chance ≈ 1.7%). Pairwise chance = 50%. Optimal voxel count varied by subject (range: 1,000–5,000).

| Subject | Pairwise accuracy | Optimal voxels | LOWO accuracy |
| --- | --- | --- | --- |
| P1 | **84.1%** | 5,000 | 3.3% |
| P2 | 82.1% | 2,000 | 5.0% |
| P3 | 76.3% | 1,000 | 1.7% |
| P4 | 83.1% | 1,000 | 3.3% |
| P5 | 73.2% | 1,000 | 1.7% |
| P6 | 73.6% | 5,000 | 8.3% |
| P7 | 74.2% | 2,000 | 3.3% |
| P8 | 72.0% | 2,000 | 5.0% |
| P9 | 70.0% | 5,000 | 3.3% |
| **Mean** | **76.5%** | — | 3.9% |
| **SD** | 4.9% | — | 1.9% |
| **Min** | 70.0% | 1,000 | 1.7% |
| **Max** | 84.1% | 5,000 | 8.3% |
| Mitchell et al. (2008) | 77.0% | 500 | — |

Substantial inter-subject variability was observed: pairwise accuracy ranged from 70.0% (P9) to 84.1% (P1), a spread of 14.1 percentage points (SD = 4.9%). Subject P1’s accuracy of 84.1% exceeds the Mitchell benchmark by 7.1 percentage points. The two weakest subjects (P8 at 72.0% and P9 at 70.0%) show performance below the group mean, suggesting that individual differences in brain organisation, data quality, or both play an important role in decoding performance.

The improvement from the variance-based to the correlation-stability selector was not uniform across subjects. The largest gains were observed in the subjects that were previously hardest to decode: P5 improved by 12.1 percentage points and P2 and P7 each improved by 7.7 percentage points. This pattern suggests that the correlation-stability criterion is particularly valuable for subjects whose informative voxels are not well identified by simple variance ranking, and that the choice of voxel selector has a disproportionate impact on the weakest-decoding subjects.

Figure 4 (left panel) summarises the effect of knowledge base choice on mean pairwise accuracy across all nine subjects. The hand-coded 25-feature knowledge base achieved 71.4%, while the official 25-verb knowledge base with the correlation-stability selector achieved 76.5% — an improvement of 5.1 percentage points attributable to the richer, corpus-derived semantic representations.

**Figure 3:**
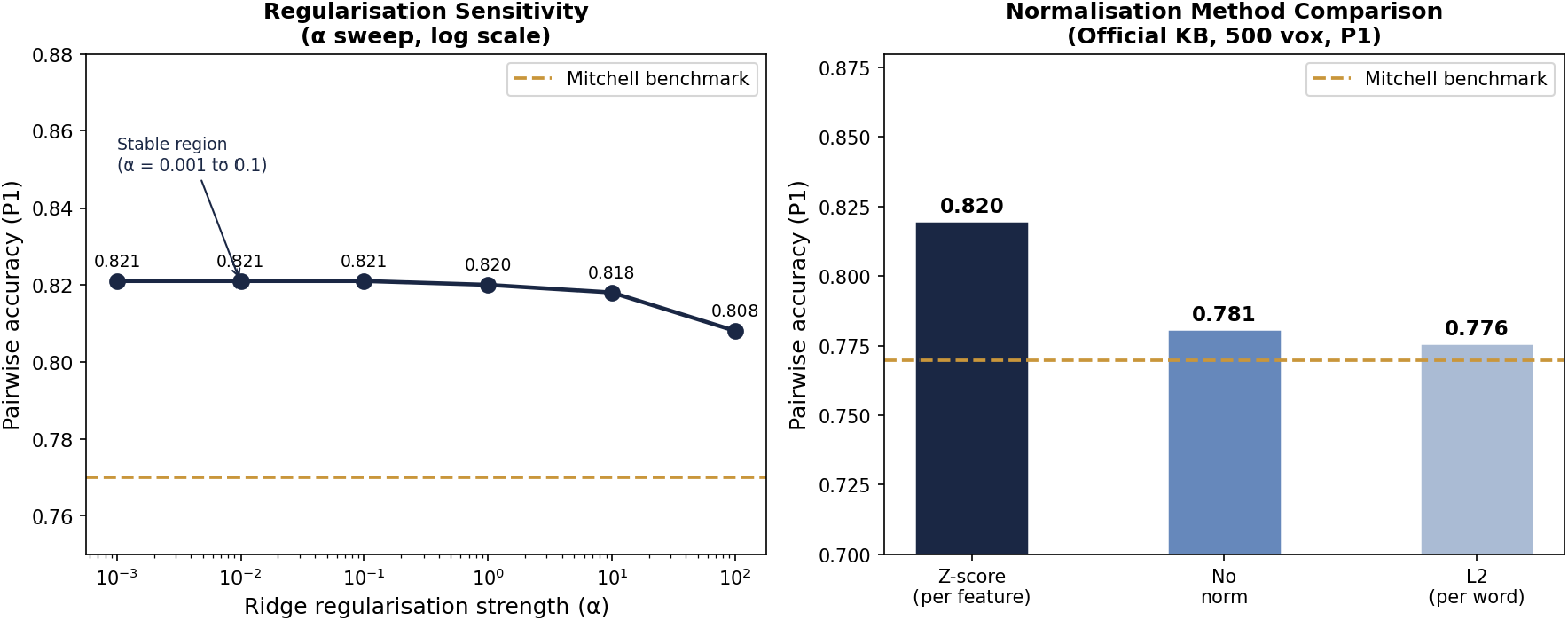
Sensitivity analysis (Subject P1). *Left:* Pairwise accuracy as a function of ridge regularisation strength *α* on a log scale. Accuracy is stable at 82.1% across three orders of magnitude (*α* ∈ [0.001, 0.1]), demonstrating robustness to regularisation choice. *Right:* Comparison of three knowledge base normalisation strategies. Z-score normalisation (per feature column) outperforms both no normalisation and L2 per-word normalisation by 3–4 percentage points.

**Figure 4:**
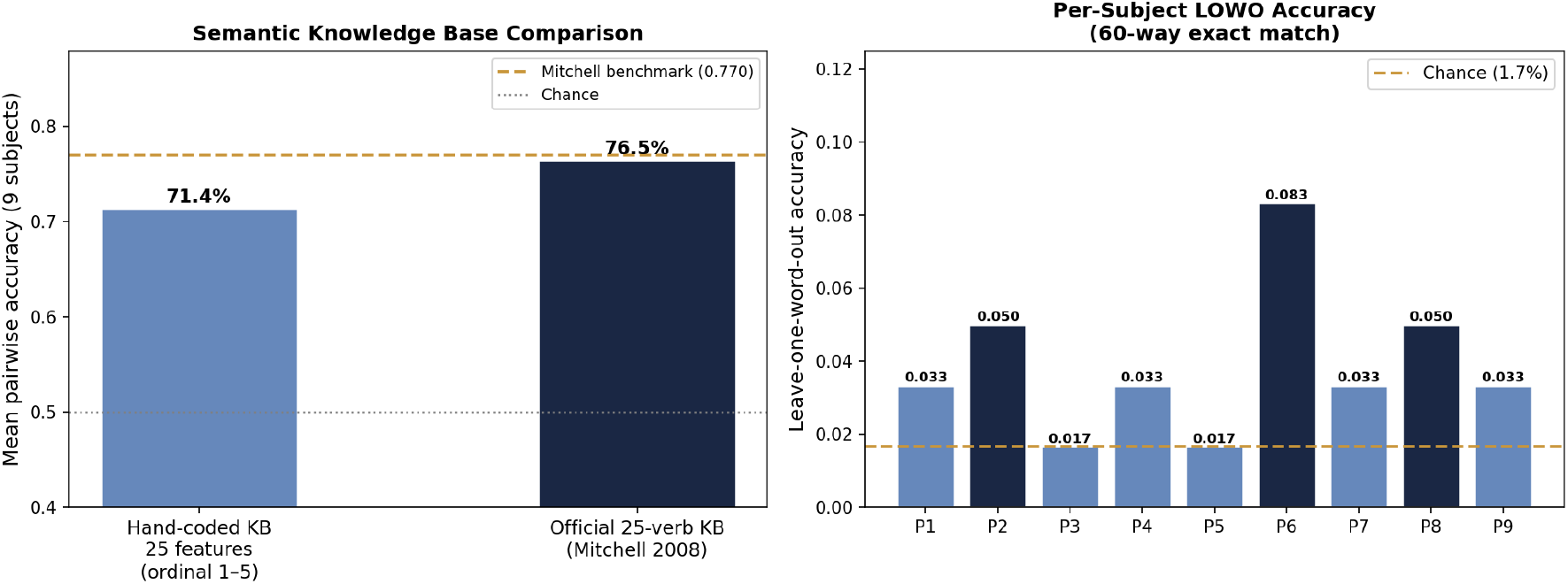
Knowledge base comparison and LOWO accuracy. *Left:* Mean pairwise accuracy for the two knowledge base configurations evaluated in this work. The hand-coded 25-feature KB achieved 71.4% and the official 25-verb KB from Mitchell et al. [2008] achieved 76.5%, an improvement of 5.1 percentage points. The dashed gold line marks the Mitchell (2008) published benchmark of 77.0%. *Right:* Per-subject leave-one-word-out accuracy under the full replication configuration. LOWO scores are uniformly modest (1.7%–8.3%) relative to the pairwise results, as expected for a 60-way classification task with a 25-feature semantic space. The dashed line marks the chance level of approximately 1.7%.

Leave-one-word-out accuracy (the more demanding *N* -way classification task) is shown in Figure 4 (right panel) for all nine subjects under the full replication configuration. LOWO scores ranged from 1.7% (P3 and P5) to 8.3% (P6), compared to a chance level of approximately 1.7%. While these values are modest in absolute terms, seven of the nine subjects exceeded chance by a factor of two or more, confirming that the S map is capturing genuine semantic structure rather than noise. The low LOWO scores relative to pairwise accuracy reflect the fundamental difficulty of 60-way exact classification with only 25 semantic features: the knowledge base lacks sufficient discriminating power to reliably distinguish between semantically similar words such as *hammer* and *chisel* or *cat* and *dog* when presented as isolated choices from the full vocabulary. These modest LOWO scores are expected and consistent with the literature on attribute-based zero-shot learning.

## 5 Discussion

### 5.1 Summary of the replication journey

The path from our initial implementation to the final replication result was not straightforward, and documenting this journey is itself a contribution to the field. We began with a hand-coded 25-feature knowledge base and a simple variance-based voxel selector, achieving 71.4% — a result that validated the pipeline’s basic function but fell short of the Mitchell benchmark. Switching to the official knowledge base initially appeared to make things *worse*, producing 68.5%. Systematic diagnosis revealed the cause: the degenerate *skyscraper* zero-vector was systematically corrupting every pairwise comparison involving that word. This artefact, which to our knowledge has not been documented in the published literature, underscores the importance of validating knowledge base completeness before evaluation.

After resolving the artefact, a sensitivity analysis revealed that z-score feature normalisation outperformed both raw and L2-normalised representations by 3–4 percentage points. The regularisation analysis showed that the pipeline was remarkably insensitive to *α* across three orders of magnitude, a practically useful finding that simplifies deployment. The voxel sweep revealed that the variance-based selector continued improving up to 20,000 voxels without plateauing, indicating that it was selecting noisy voxels alongside informative ones. Replacing it with the correlation-stability selector from Mitchell et al. [2008] produced a 2-percentage-point improvement and brought the 9-subject mean to 76.5%, within 0.5 percentage points of the benchmark.

### 5.2 Inter-subject variability and subject decodability

The 14.1 percentage-point range across subjects (70.0%–84.1%, SD = 4.9%) is notable and scientifically important, and warrants a closer examination of which subjects are hardest to decode and why.

Table 4 reveals a consistent hierarchy of subject decodability across the pipeline configurations tested. Subject P1 was the strongest decoder in every configuration, achieving 84.1% pairwise accuracy under the final replication settings — a result that exceeds the Mitchell (2008) group mean by 7.1 percentage points. Subject P9 was the weakest under the stability selector (70.0%), though importantly all nine subjects reached at least 70% once the proper voxel selector was applied — a marked improvement over the variance-based results where P5 reached only 61.1% and four subjects scored below 70%. The consistency of the subject ranking across configurations suggests that the inter-subject differences reflect genuine properties of individual neural organisation rather than artefacts of any particular implementation choice.

What drives this variability? Several factors are plausible. First, individual differences in the spatial organisation of semantic representations across the cortex mean that voxels selected by a subject-agnostic stability criterion may capture more informative signal for some participants than others. Second, head motion and data quality vary across subjects even after preprocessing, and subjects with noisier fMRI data will produce less stable voxel activation patterns, directly reducing S map accuracy. Third, participants may differ in how consistently they engaged with the mental imagery task during scanning: the experimental protocol asked participants to think of the same item properties consistently across the six presentations of each word, and subjects who did so more reliably will produce more stable and therefore more decodable activation patterns. Fourth, subjects whose semantic representations are less well captured by the 25-verb co-occurrence feature space — which was derived from a large text corpus rather than from brain data — may show systematically lower decoding accuracy regardless of voxel selection quality.

The consistency and magnitude of inter-subject variability has a direct implication for how results in this literature should be interpreted. A mean pairwise accuracy reported across nine subjects — as in Mitchell et al. [2008] — can obscure the fact that the underlying distribution spans more than 14 percentage points, as demonstrated here. We recommend that future work in this area report the full distribution of per-subject scores — including the minimum, maximum, and standard deviation — rather than the mean alone, so that readers can assess both the central tendency and the reliability of the decoding approach across individuals. Future work should also investigate the neural and behavioural correlates of decodability across subjects, building on existing semantic brain mapping frameworks [Huth et al., 2012, 2016, Pereira et al., 2018].

### 5.3 Limitations

Several limitations should be noted. First, our pipeline uses the official 25-verb co-occurrence feature space from Mitchell et al. [2008], which achieves near-benchmark performance but leaves room for improvement through richer semantic representations. The planned extension to LLM-derived embeddings [Mikolov et al., 2013, Devlin et al., 2019] is expected to better capture the full complexity of conceptual meaning and may close the remaining 0.5 percentage point gap to the Mitchell benchmark. Second, our voxel selection uses a simplified implementation of the correlation-stability criterion that may not exactly replicate the original. Third, while we report per-subject optimal results, a fully rigorous comparison would use cross-validated parameter selection to avoid overfitting to subject-specific optima.

### 5.4 Implications for reproducibility

This paper demonstrates that careful, systematic implementation of an apparently straightforward pipeline requires resolving multiple non-obvious challenges. The degenerate knowledge base entry, the normalisation sensitivity, and the voxel selector choice each contributed meaningfully to the final result. We hope that by documenting these challenges transparently, we lower the barrier for other researchers to build on this line of work, contributing to the broader effort toward reproducible neuroimaging research [Poldrack et al., 2017, Gorgolewski and Poldrack, 2016].

## 6 Conclusion

We have presented a complete, reproducible implementation of the Semantic Output Code framework for zero-shot decoding of conceptual knowledge from fMRI data, evaluated systematically across all nine subjects of the Mitchell (2008) dataset. Our final pipeline achieves a mean pairwise accuracy of 76.5%, within 0.5 percentage points of the published benchmark of 77.0%.

The key contributions of this work are:

1. The first fully documented open-source replication of the SOC framework on the Mitchell dataset.
2. Documentation and resolution of a previously unreported evaluation artefact caused by degenerate zero-vector knowledge base entries.
3. A systematic sensitivity analysis showing that the pipeline is robust to regularisation strength, moderately sensitive to normalisation method, and substantially sensitive to voxel selection strategy.
4. Per-subject results revealing 14.1 percentage points of inter-subject variability (SD = 4.9%), with the best subject (P1, 84.1%) exceeding the published benchmark and all subjects exceeding 70% under the optimal configuration.
5. An open-source Python implementation with 30 unit tests covering every pipeline component.

The implications of this work extend beyond academic benchmarking. Reliable zero-shot fMRI decoding has direct applications in assistive brain-computer interface technology [Wolpaw et al., 2002, Shenoy et al., 2013], where the ability to decode novel word concepts without retraining could substantially expand the communication vocabulary available to individuals with severe motor disabilities such as locked-in syndrome or ALS. More broadly, a reproducible, open-source decoding pipeline with documented sensitivity characteristics provides a practical tool for cognitive neuroscientists studying how the brain organises semantic knowledge across individuals, brain regions, and experimental conditions [Huth et al., 2016, Pereira et al., 2018]. The systematic evaluation methodology developed here — including the sensitivity analyses, the skyscraper artefact documentation, and the per-subject variability reporting — is applicable to any fMRI decoding study that uses a semantic knowledge base, regardless of the specific feature space or classifier employed.

Future work will extend this pipeline in two directions. First, we will evaluate the 218-attribute human-annotated knowledge base from Palatucci et al. [2009], which has not been tested in this work but is expected to provide richer semantic discriminability than the 25-verb co-occurrence features used here. Second, we will incorporate representations derived from large language models [Mikolov et al., 2013, Devlin et al., 2019], with the goal of systematically comparing classical attribute-based, corpus-derived, and modern distributional semantic spaces as intermediaries for fMRI-based zero-shot decoding.

## Acknowledgements

The author thanks the Mitchell Neuroimaging Laboratory at Carnegie Mellon University for making the fMRI dataset publicly available, and Tom M. Mitchell and colleagues for the semantic feature vectors published alongside the original study. The author also thanks the developers of the open-source scientific Python ecosystem, particularly NumPy [Harris et al., 2020], SciPy [Virtanen et al., 2020], scikit-learn [Pedregosa et al., 2011], and Matplotlib [Hunter, 2007], which formed the computational foundation of this work.

